# Investigating Sensitivity, Specificity and Accuracy of Variant Calling Pipelines for Analyzing SARS-CoV-2 Data

**DOI:** 10.1101/2024.01.24.576385

**Authors:** Aadi C. Krishna, Judy S. Choi

**Affiliations:** Shiv Nadar School Noida, India; Korea International School, South Korea

## Abstract

The rapidly increasing popularity of Next Generation Sequencing and analysis methods in clinical and research settings necessitates an understanding of ideal combinations in identifying genomic variants. Especially with the importance of detecting accurate variants for the development of targeted SARS-CoV-2 vaccines. This research compares the results of two ‘Mapping Algorithms ‘, BWA-MEM and Bowtie2, and two ‘Variant Calling Algorithms ‘, LoFreq and FreeBayes, and their combinatory Variant Calling Pipelines on the analyses of Next Generation Sequencing (NGS) data of five SARS-CoV-2 samples collected from patients in the USA, India, Italy, and Malawi and sourced for this research from the publicly available NCBI SRA database. Our analysis of mapping algorithms found that BWA-MEM likely has higher sensitivity and specificity than Bowtie2 for mapping reads, and their specificity and sensitivity vary with read length. Furthermore, the accuracy of variant calling algorithms increases with the number of reads, while higher read length possibly leads to divergence in accuracy and sensitivity. Overall, FreeBayes was found to likely be more sensitive to detecting variants when used with Bowtie2 rather than BWA-MEM for analyzing SARS-CoV-2 data.

## I. Introduction

A Variant Calling Pipeline is the process of identifying variants in a genomic sequence data sample using one mapping and variant calling algorithm [1]. Mapping using a mapping algorithm is the first step in the Variant Calling Pipeline to map the raw reads collected by the NGS sequencer to the reference, in our case, the SARS-CoV-2 genomic data. The reads identified as unmapped and not properly paired are eliminated from further analysis. A variant calling algorithm is used in the final step of the variant analysis to determine the mutations, Single Nucleotide Variations (SNV) and Indels, in the sample and their locations. Hence, the results of mapping algorithms and variant calling algorithms directly influence the outcome of studies, and it is crucial to investigate the characteristics of mapping algorithms and variant calling algorithms that deviate with the nature of the genome to effectively design Variant Calling Pipelines to understand variants of SARS-CoV-2 best and develop successful regional vaccines specifically tailored to the mutations.

Our research focuses on 5 SARS-CoV-2 samples from four countries to analyze the sensitivity, specificity, and accuracy of the mapping algorithms and variant calling algorithms, and their combinatory Variant Calling Pipelines. Sensitivity is defined as the ability to correctly identify the positive results, and specificity is the ability to correctly identify the negative results. For the research, we chose two of the most commonly used mapping algorithms, BWA-MEM and Bowtie2, and variant calling algorithms, LoFreq and FreeBayes [2].

## II. Methods

The National Center for Biotechnology Information ‘s SRA Run Selector database was utilized to select five SARS-CoV-2 (2019-nCoV) coronavirus samples: two from USA, and one each from India, Italy and Malawi. The UseGalaxy [3] project ‘s tools were employed to analyze the collected sequence data. Firstly, each raw FASTQ file, which stores the nucleotide sequence for a sample, was “De-interlaced” to split the paired-end sequencing data into two, each containing one of the reads and its base quality scores. Both read files were made to undergo quality checks and generate initial statistics using FASTQC.

The reference SARS-CoV-2 genome (NC_045512.2) was collected from the NCBI website. Then, BWA-MEM and Bowtie2 were employed to map the raw reads to the reference and generate a Binary Alignment Map (BAM) that is the raw alignment data file of the genome sequence Following the mapping, Samtools Stats was used to generate detailed statistics from the BAM files: identification of “Mapped Reads,” “Not Properly Paired Reads,” etc. Next, “Unmapped” and “Not Properly Paired Reads” were eliminated using the Filter BAM tool. Additionally, all reads with a mapping quality less than 20 Phred score were also filtered to improve accuracy.

Next, variant detection was performed with LoFreq and FreeBayes on the filtered mapped file, generating four Variant Calling Files (VCF) with all the combinations of the variant calling algorithms and mapping algorithms for each sample. Each file enumerated the SNVs and Indels in the samples compared to the reference using one mapping algorithm and variant calling algorithm. Lastly, the four VCF files – BWA-MEM + LoFreq; Bowtie2 + LoFreq; BWA-MEM + FreeBayes; and BWA-MEM + FreeBayes, were intersected to end up with 1 VCF file for each sample.

## III. RESULTS AND DISCUSSION

Table I details the initial statistics - read length and total reads collected for all the samples from the FASTQC files.

**TABLE I.** Initial Statistics of the 5 Selected Samples.

| Genome | Country | Read Length (bp) | Total Reads |
| --- | --- | --- | --- |
| SRR13684562 | USA (1) | 251 | 126274 |
| SRR14671720 | USA (2) | 75 | 716228 |
| SRR14777079 | Italy | 36 | 5117166 |
| SRR14877919 | India | 149 | 87878 |
| SRR14698004 | Malawi | 251 | 416276 |

The results of the subsequent mapping algorithms and filtering are presented in Table II. “Properly Paired Reads” found by BWA-MEM were more for 3 out of the 5 samples, with only the USA (1) and India samples giving a 4.9% and 0.35% higher result for Bowtie2. Furthermore, BWA-MEM consistently found a larger number of “Mapped Reads” than Bowtie2, with an average 2.5% higher result. After filtering, the total remaining reads were on average 2.94% greater in BWA-MEM for three samples, while Bowtie2 returned a higher number for the USA and India samples.

**TABLE II.** Result Statistics of Bwa-mem and Bowtie2 Mapping.

| Sample | Mapping | Properly Paired Reads | Mapped Reads | Remaining Reads After Filtering |
| --- | --- | --- | --- | --- |
| USA (1) | BWA- MEM | 111,446 | 119,156 | 111,442 |
|  | Bowtie2 | 116,926 | 118,291 | 116,176 |
| USA (2) | BWA- MEM | 648,460 | 668,461 | 645,504 |
|  | Bowtie2 | 616,176 | 636,437 | 615,654 |
| Italy | BWA- MEM | 4,670,076 | 4,714,133 | 4,670,001 |
|  | Bowtie2 | 4,628,812 | 4,695,130 | 4,627,114 |
| India | BWA- MEM | 78,490 | 83,864 | 78,488 |
|  | Bowtie2 | 78,760 | 80,421 | 78,642 |
| Malawi | BWA- MEM | 343,452 | 343,841 | 343,219 |
|  | Bowtie2 | 334,256 | 339,071 | 331,954 |

The output of variant calling algorithms, LoFreq and FreeBayes, are presented in Table III. LoFreq returned more variants in the USA (1), USA (2), and Italy samples; however, FreeBayes found more variants in the India and Malawi samples. USA (1), India, and Malawi showed a difference of more than two times between LoFreq and FreeBayes results; whereas USA (1) and Malawi were the only two samples with lower values for LoFreq with Bowtie2 compared to LoFreq with BWA-MEM. FreeBayes with BWA-MEM consistently found fewer or equal number of variants as compared to FreeBayes with Bowtie2. LoFreq and FreeBayes found the most similar results of variants in the Italy sample, with a 4.75σ standard deviation between the variants found by variant calling algorithms and the final intersected file, followed by the USA (2) with 15.05σ.

**TABLE III.** Number of Variants Found By Each Variant Calling Pipeline.

| Sample | LoFreq |  | FreeBayes |  | Final Intersected |
| --- | --- | --- | --- | --- | --- |
|  | BWA-MEM | Bowtie 2 | BWA-MEM | Bowtie 2 |  |
| USA (1) | 327 | 305 | 25 | 25 | 14 |
| USA (2) | 50 | 75 | 46 | 53 | 28 |
| Italy | 41 | 45 | 38 | 42 | 31 |
| India | 32 | 35 | 61 | 65 | 24 |
| Malawi | 40 | 39 | 93 | 104 | 25 |

## IV. Conclusion

The study focused on analyzing 5 samples from four different countries around the world, mapping the reads collected by previous studies to the COVID-19 reference genome, and finding the variants in each sample. Through this process, two mapping algorithms, BWA-MEM and Bowtie2, and two variant calling algorithms, LoFreq and FreeBayes, were evaluated.

The greater number of “Mapped Reads” found by BWA-MEM shows its likely higher sensitivity and specificity in mapping reads to the reference compared to Bowtie2. The larger values of “Properly Paired Reads” only in the USA (1) and India samples could be a result of two possibilities: the lower number of reads present in the two samples and poorer read quality, or a variation in the specificity and sensitivity of Bowtie2 and BWA-MEM with an increasing number of reads in the sample, consistent with the findings of [4], since USA (1) and India had the lowest number of total reads.

All samples showed more variants detected by Bowtie2 + FreeBayes, except USA (1) that found the same variants as BWA-MEM + FreeBayes. Thus, it is believed that FreeBayes is likely to be more sensitive to finding variants when Bowtie2 is used as the mapping algorithm. Fewer variants found by Bowtie2 + LoFreq only in the USA (1) and Malawi samples can possibly be explained by their longer sequence length, 251, implying that the sensitivity and specificity of LoFreq changes with sequence length [5]. Furthermore, there appears to be an effect of the number of reads, depth of reads and sequence length on the significant differences in variants identified by FreeBayes and LoFreq algorithms in the samples. The accuracy, measured using the similarity in the number of variants identified by both the variant calling algorithms, increases with the number of reads and depth of reads, and decreases with higher sequence length, as demonstrated by the Italy and USA (2) samples. However, besides the aforementioned relationships, based on previous studies [6], it can be reasoned that the coverage of the samples, which was not calculated for analysis in this research, may also be a factor behind the varying sensitivity of the two algorithms.

Therefore, it is recommended that for an analysis of SARS-CoV-2 genome sample, the read length, number of reads, read depth, and coverage of the sample are collected, the trends discussed here are considered, and accordingly the ideal mapping algorithm and variant calling algorithm combination for the Variant Calling Pipeline is decided. Additionally, it is suggested to collect NGS SARS-CoV-2 data with shorter read lengths. Further researchers may extend this work and explore the sensitivity, specificity, and accuracy of other Variant Calling Pipelines for analyzing SARS-CoV-2 data through alternate mapping algorithms – Novoalign – and variant calling algorithms – DeepVariant.

## Acknowledgement

This study was conducted as part of the International Science Engagement Challenge (ISEC, summeratisec.org). We would like to thank our mentor, Elif Bozlak, for her guidance.

